# Brain macrophages acquire distinct transcriptomes prior to demyelination in multiple sclerosis

**DOI:** 10.1101/2021.10.27.465877

**Authors:** Anneke Miedema, Emma Gerrits, Nieske Brouwer, Qiong Jiang, Laura Kracht, Michel Meijer, Erik Nutma, Regina Peferoen-Baert, Anna T.E. Pijnacker, Evelyn M. Wesseling, Marion H.C. Wijering, Hans-Joachim Gabius, Sandra Amor, Bart J.L. Eggen, Susanne M. Kooistra

## Abstract

Multiple sclerosis (MS) is a disease of the central nervous system (CNS) that is characterized by inflammation and focal areas of demyelination, ultimately resulting in axonal degradation and neuronal loss. Several lines of evidence point towards a role for microglia and other brain macrophages in disease initiation and progression, but exactly how lesion formation is triggered is currently unknown. Here, we characterized early changes in MS brain tissue through transcriptomic analysis of normal appearing white matter (NAWM). We found that NAWM was characterized by enriched expression of genes associated with inflammation and cellular stress derived from brain macrophages. Single cell RNA sequencing confirmed an early stress response in brain macrophages in NAWM and identified specific macrophage subsets that associate with different stages of demyelinating lesions. These early changes associated with lesion development in MS brain tissue may provide therapeutic targets to limit lesion progression and demyelination.

## Introduction

Multiple sclerosis (MS) is an auto-immune disease damaging the central nervous system (CNS) and affects 2.8 million people worldwide[42]. MS is characterized by demyelinated lesions in the brain, optic nerves and spinal cord, ultimately resulting in damage to the axons which leads to cognitive problems, blindness, impaired motor function or even paralysis, depending on the location and extent of the lesions[5]. Despite extensive efforts, the factors causing disease and initiating the formation of new lesions are not yet known. Genome wide association studies (GWAS) point towards a role of microglia, the resident macrophages in the parenchyma of the brain, in MS pathology[15]. Besides microglia, other types of macrophages have shown to be affected in mouse models for MS, for example perivascular macrophages, meningeal macrophages, choroid plexus macrophages and infiltrating peripheral monocytes, collectively termed CNS-associated macrophages (CAMs)[16].

Additional clues towards the processes involved in lesion initiation are derived from the analysis of the non-lesioned areas from the CNS of affected individuals, such as normal-appearing white matter (NAWM). Compared to healthy controls, NAWM shows several pathological abnormalities[1, 25]. For example, perturbed myelin-axon interactions have been described as an early event in MS NAWM[25]. Additionally, morphologically altered and HLA-DR-expressing macrophages are detected in NAWM[1]. Analysis of gene expression levels indicated that inflammatory processes are already ongoing in NAWM and might involve both microglia and astrocytes[12, 28, 32, 37, 40]. However, the underlying pathological mechanisms initiating and driving (inflammatory) demyelination in MS remain incompletely understood, and likely involve multiple cell types both within and outside the CNS[6]. To identify early changes in MS brain tissue, we here analyzed the transcriptional profile of NAWM tissue without apparent macrophage activation (in situ) and applied single-cell RNA sequencing to further determine changes in brain macrophages, including microglia.

## Results

### Transcriptomic changes in normal appearing white matter

To identify global disease-associated transcriptomic changes in the brains of MS donors, total tissue transcriptomic profiling was performed on white matter from control donors (CWM), normal appearing white matter (NAWM) from MS donors and demyelinated white matter lesions (WMLs) from MS donors (Fig. 1A, Supplementary Table S1). To avoid the presence of demyelinated lesions or overt activation of macrophages in NAWM samples, tissue sections immediately adjacent to the tissue sections used for RNAseq were stained for PLP1 and HLA-DR, where for NAWM the PLP1 score must be 10 to indicate intact myelin and HLA-DR score < 4 representing low-grade macrophage activation status (Fig. 1A, Supplementary Fig. S1A, Supplementary table S1). Principal component analysis segregated the WML samples from CWM and NAWM samples in the first principal component (PC1). Furthermore, a clear segregation between NAWM and CWM samples was observed in PC2 and some NAWM samples grouped with WML samples, indicating that the NAWM in MS brains is affected prior to demyelination (Fig. 1B). Differential gene expression analysis confirmed the differences between the groups by identification of hundreds of differentially expressed genes (DEGs; abs(logFC) > 1 and FDR < 0.05) between NAWM vs CWM samples, WML vs NAWM and WML vs CWM (Fig. 1C, Supplementary Table S2). Biological processes associated with DEGs enriched in NAWM compared to CWM indicate that apoptosis, stress and inflammatory responses are present in NAWM tissue prior to lesion formation (Fig. 1D). Additionally, genes reported as immediate-early genes (IEGs), reported to be the first genes being transcribed when cells are stimulated such as during an inflammatory response, were enriched in NAWM compared to CWM (Supplementary Fig. S1B, C). DEGs uniquely enriched in WML compared to NAWM were associated with epithelial cell apoptosis and cilium assembly and movement, indicating that these processes occur during or after demyelination/lesion formation. DEGs associated with extracellular matrix organization, humoral immune response, leukocyte migration and complement activation were enriched in both NAWM and WML samples compared to CWM (Fig. 1D).

**Figure 1.**
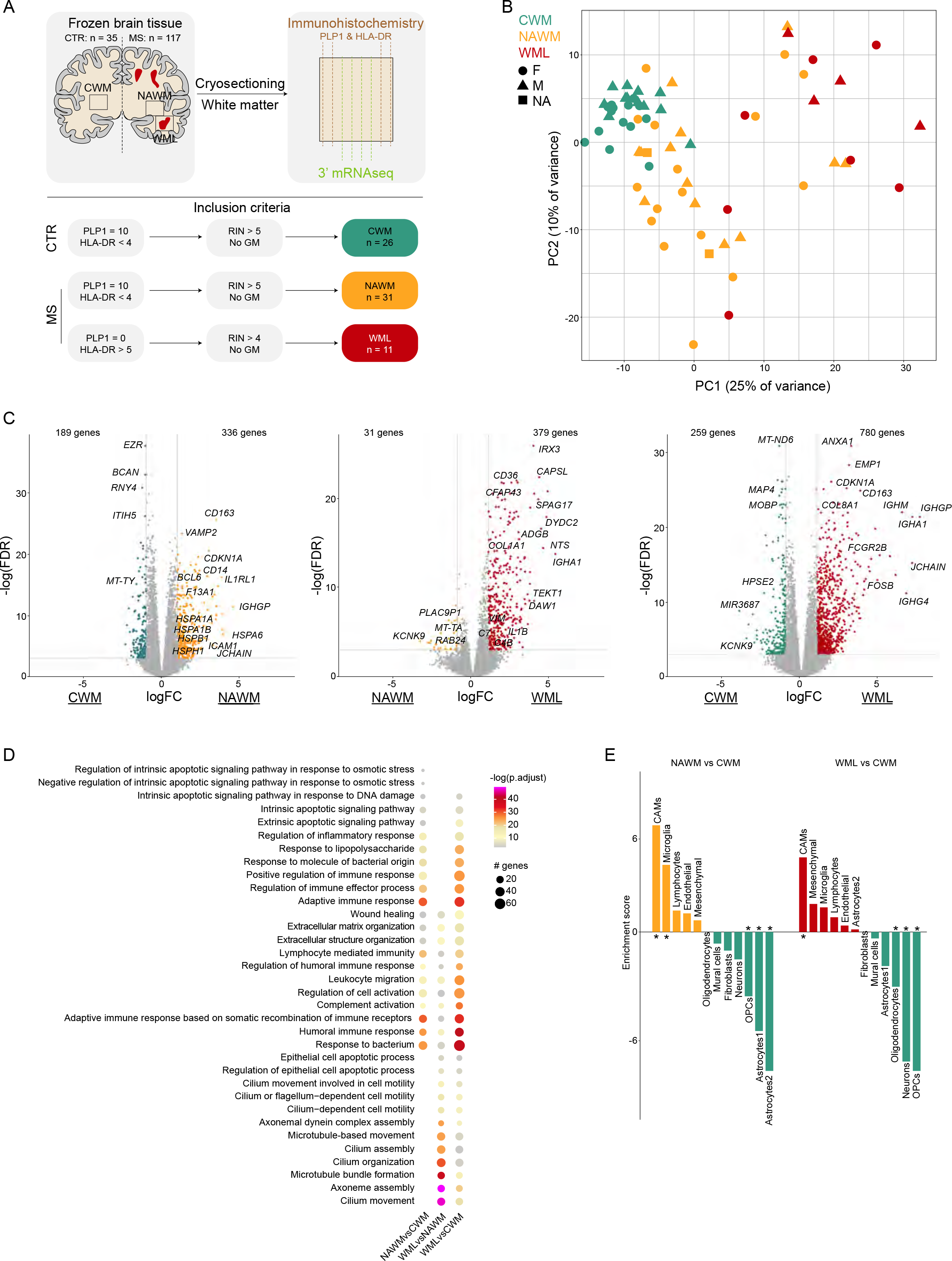
Transcriptomic changes in normal appearing white matter. (**A**) Schematic overview of the experiment. (**B**) Principal component analysis of 68 white matter brain tissue samples. Colors depict donor groups. Shapes depict sex. (**C**) Volcano plots depicting differential gene expression between NAWM vs CWM samples (left), WML vs NAWM samples (center) and WML vs CWM samples (right). Colored symbols indicate significantly differentially expressed genes (logFC > 1 and adjusted p-value < 0.05). (**D**) Dot plot depicting the 10 most significant gene ontology analyses associated with DEGs from the comparisons indicated on top. Size depicts # of genes in gene ontology set, color scale depicts significance level. (**E**) Bar plot depicting expression-weighted gene set enrichment analysis of DEGs between NAWM versus CWM (left) and WML vs CWM (right) based on cell type specific gene signatures derived from human snRNAseq study (Gerrits et al. 2021). *: p <0.05; Abbreviations: CWM = control white matter (CTR donors); NAWM = normal appearing white matter (MS donors); WML = white matter lesion (MS donors).

To elucidate whether the identified DEGs are caused by changes in cell type composition we performed expression-weighted cell type enrichment analysis (EWCE)[34]. DEGs enriched in NAWM when compared to CWM were significantly enriched in CAMs and microglia and depleted in neurons, oligodendrocyte precursor cells (OPC) and astrocytes. DEGs between WML and CWM were significantly enriched for CNS-associated macrophage (CAM)-genes and depleted for oligodendrocyte-, neuron- and OPC-genes (Fig. 1E). This suggests that these cell types have a different abundance and/or disease-associated transcriptomic changes in gene expression profiles between the conditions.

### Inflammation and stress gene modules are enriched in normal appearing white matter

To extract process- or cell type-specific gene modules a weighted gene co-expression network analysis (WGCNA) was performed (Fig. 2A, Supplementary Table S3), and combined with EWCE to identify whether modules contained cell type specific gene expression profiles (Fig. 2B). The majority of the WGCNA-identified gene modules were enriched in oligodendrocytes, indicating that these cells are amongst the most affected cell types in MS brains, in line with the known pathological features of MS lesions (Fig. 2B). However, not all oligodendrocyte-associated modules were depleted in WML compared to CWM and NAWM (‘red’, ‘green’, ‘brown’, ‘purple’, ‘cyan’, ‘yellow’; Supplementary Fig. S1D). This indicates that transcriptomic changes also occur within the oligodendrocyte population, in the absence of loss of oligodendrocyte numbers in WMLs, or that specific subpopulations of oligodendrocytes are lost in WMLs. Additionally, many oligodendrocyte modules were differentially abundant between CWM and NAWM, where no demyelination was present (‘red’, ‘green’, ‘brown’, ‘tan’, ‘cyan’, ‘yellow’; Supplementary Fig. S1D). One such module, ‘tan’, was enriched in NAWM compared to CWM (Fig. 2C) and comprised both oligodendrocyte and astrocyte genes (Fig. 2B). Module ‘tan’ was associated with ‘stress response’, ‘metabolism’ and ‘cell cycle’, suggesting that prior to demyelination oligodendrocytes and/or astrocytes display an early stress response (Fig. 2C). Two gene modules that were enriched in astrocytes were detected (‘black’ and ‘midnightblue’) and both were depleted in NAWM samples compared to CWM (Fig. 2D, Supplementary Fig. S1D). Both modules contained genes associated with typical astrocyte functions, indicating that astrocytic support may be depleted in NAWM tissue (Fig. 2D).

**Figure 2.**
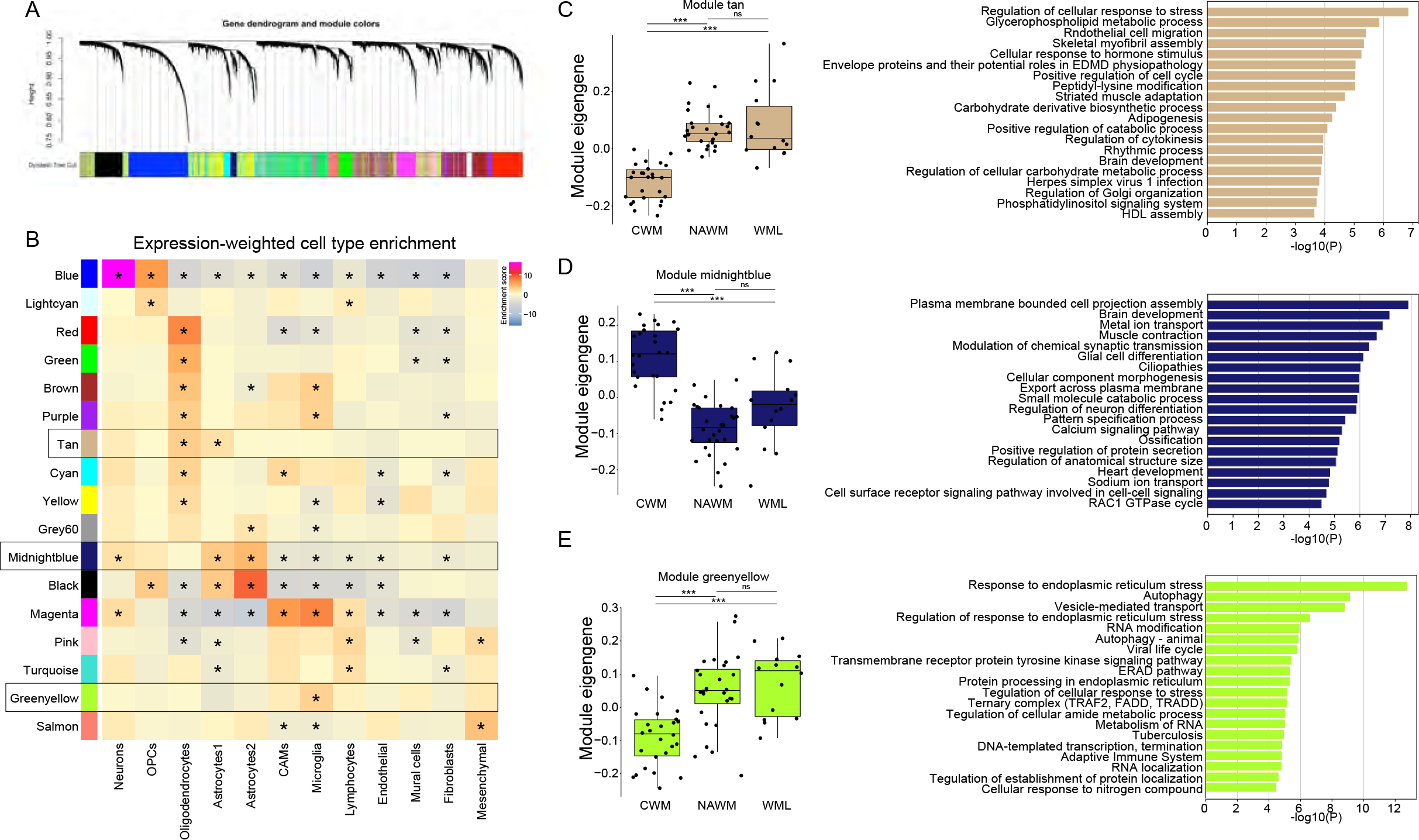
Inflammation and stress gene modules are enriched in normal appearing white matter. (**A**) Gene dendrogram and module colors of weighted gene correlation network analysis. (**B**) Heatmap depicting enrichment scores per module of expression-weighted cell type enrichment analysis. (**C-E**) Boxplot depicting module eigen genes for the tan, midnightblue and greenyellow gene modules (left). Bar plot depicting gene ontology analysis of tan, midnightblue and greenyellow gene modules (right). *: p <0.01; **: p < 0.001; ***: p < 0.0001. Abbreviations: CWM = control white matter (CTR donors); NAWM = normal appearing white matter (MS donors); WML = white matter lesion (MS donors).

Module ‘magenta’, significantly enriched in WML samples, comprised genes associated with inflammation, and was associated with CAMs, microglia and lymphocytes, indicating that the abundance of these cells is increased in WML or that they shift towards a more pro-inflammatory profile (Supplementary Fig. S1D, E, Supplementary Table S3). Module ‘pink’ also comprised genes associated with inflammation, and this module was particularly enriched in lymphocytes. Interestingly, module ‘pink’ was also enriched in NAWM vs CWM samples, suggesting lymphocyte infiltration in the brain occurs prior to demyelination (Supplementary Fig. S1D, E, Supplementary Table S3).

Module ‘greenyellow’ was enriched in microglia (Fig. 2B) and was more abundant in NAWM and WML samples compared to CWM. This module was comprised of genes associated with stress response and autophagy, indicative of cellular stress in microglia prior to demyelination (Fig. 2E). These data suggest that the increase in inflammation-, stress- and apoptosis-related gene expression in NAWM and WML total tissue compared to CWM (Fig. 1C, D) are derived from both oligodendrocytes and brain macrophages in NAWM prior to demyelination.

### Transcriptomic heterogeneity of macrophages in MS brain tissue

Both stress- and inflammation-related gene expression patterns were associated with brain macrophages in NAWM tissue. Therefore, to investigate whether the detected expression changes were due to altered gene expression patterns within specific macrophage subpopulations in MS brains, single-cell RNAseq was performed on CD45^pos^CD11B^pos^ macrophages isolated from NAWM tissue and tissue with macroscopically visible demyelinated white matter (Fig. 3A). Normal-appearing tissue from the cortex from the same donors was included to obtain sufficient cells with contrasting signatures to identify disease-associated signatures and heterogeneity in NAWM and WML samples whilst avoiding donor variation.

**Figure 3.**
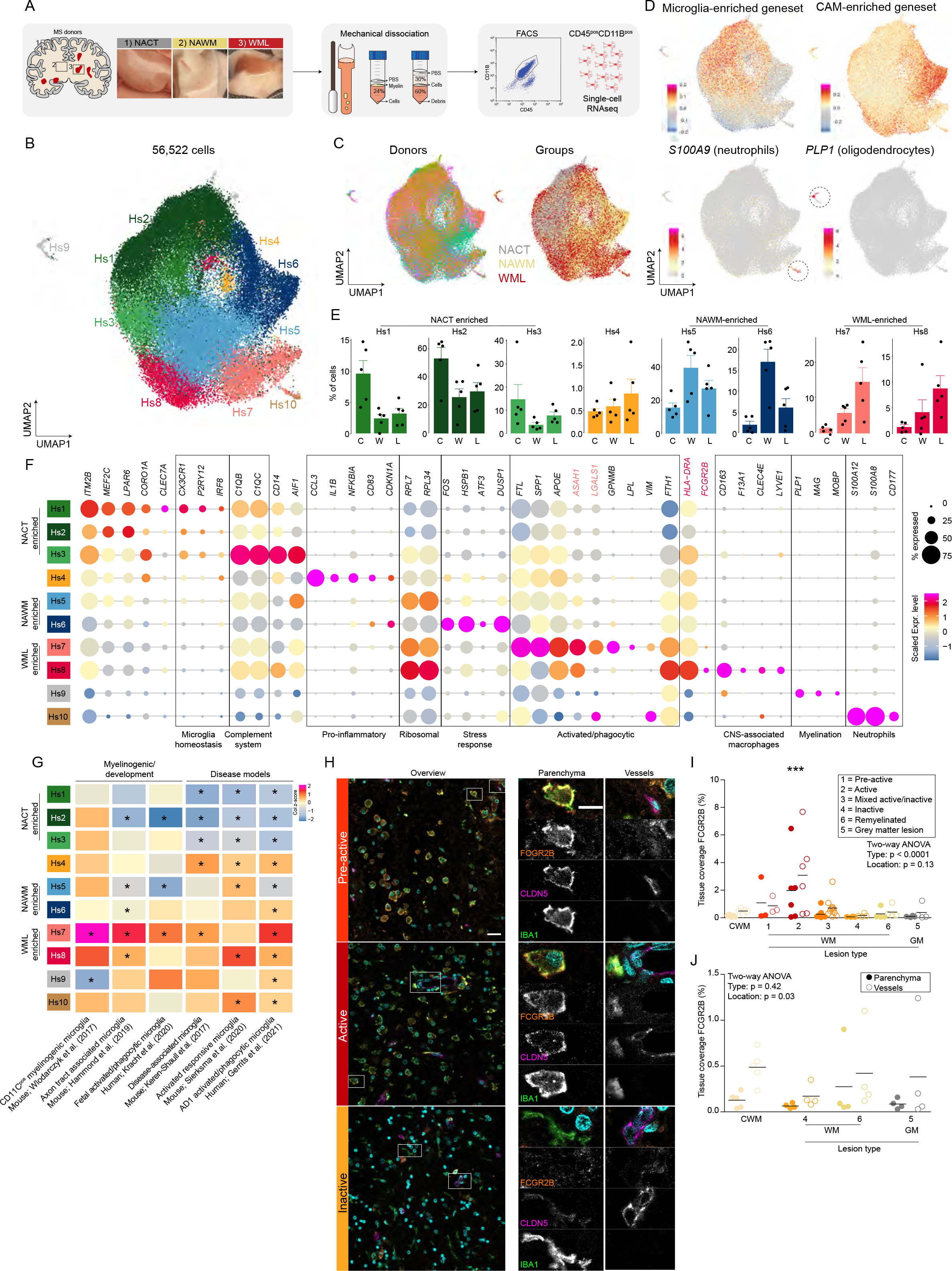
Brain macrophage heterogeneity in multiple sclerosis brain tissue. (**A**) Workflow of the experiment. (**B**) UMAP depicting 56,522 cells. Colors depict unsupervised clustering. (**C**) UMAP where colors depict donor of origin (left) or tissue region of origin (right). (**D**) UMAPs where color scales depict module scores of microglia- and CAM-enriched gene sets (top) or expression levels of *S100A9* and *PLP1* (bottom). (**E**) Bar plots depicting cluster distribution among donor groups with standard error. Symbols depict individual samples. C = NACT; W = NAWM; L = WML. (**F**) Dot plot depicting marker genes of each cluster. Size of the symbols depicts the fraction of cells expressing the gene, color scale depicts average expression level. (**G**) Heatmap depicting enrichment scores of literature-derived gene-sets in the scRNAseq dataset. (**H**) Immunofluorescent images of stainings for FCGR2B (orange; Hs8); CLDN5 (magenta; blood vessels), Hoechst (cyan; nuclei) and IBA1 (green; macrophages) in a pre-active, active and inactive lesion. Left panels: overview image. Scale bar = 25 μm. Center panel: Zoom in on parenchymal macrophages. Scale bar = 10 μm. Right panel: Zoom in on vessel-associated macrophages. (**I**) Quantification of immunohistochemistry for FCGR2B in parenchyma and near/on blood vessels. *: p <0.05. Abbreviations: NACT = normal appearing cortical tissue; NAWM = normal appearing white matter; WML = white matter lesion.

Macrophages were isolated from fresh post-mortem brain tissue within 12 hours after autopsy using our mechanical isolation protocol for human microglia[9], performing the entire procedure at 4°C to minimize possible *ex vivo* activation of cells (Fig. 3A, Supplementary Fig. S2A). In total, 56,522 CD45^pos^CD11B^pos^ cells were analyzed using single-cell RNA sequencing (scRNAseq; Supplementary Fig. S2A, B) and these were grouped into 10 subtypes (Hs 1-10) using unsupervised clustering (Fig. 3B). None of the clusters was donor-specific, but a segregation based on tissue origin was observed where particularly NACT and NAWM derived cells segregated in the UMAP (Fig. 3C, Supplementary Fig. S2C). Significant differences in gene expression between NACT and NAWM-derived cells were observed, where cortical tissue derived cells were associated with phagocytosis and synapse pruning and white matter macrophages with stress and apoptosis (Supplementary Fig. S2C-E, Supplementary Table S5). The expression of oligodendrocyte and neutrophil genes were enriched in two small clusters (Hs9 and Hs10) and likely represent a small number of non-macrophage cell types derived from the CNS parenchyma and the small amount of blood present in the vessels at the time of autopsy (Fig. 3D, Supplementary Table S4).

Recently, we reported the distinct transcriptomic profiles of microglia and other CNS-associated macrophages (CAMs) in our single-nucleus RNA sequencing (snRNAseq) dataset of human cortical brain tissue[11]. To distinguish between different types of brain macrophages in the present dataset, gene set module scoring was performed using marker genes specific for microglia or CAMs.

Expression of the CAM marker geneset was enriched in clusters Hs6 and Hs8 (Fig. 3D), and expression of typical CAM genes (*CD163*, *F13A1*, *LYVE1;* [16]) was enriched in cluster Hs8 and these were most abundant in WML samples (Fig. 3E, F). Cluster Hs6 was particularly enriched in the NAWM samples, whereas cluster Hs8 was enriched in WML samples (Fig. 3E). The identified gene signatures for the human macrophage subtypes we identified was comparable with macrophage/microglia subsets identified in other studies (Fig. 3G). In MS, distinct lesion types can be identified that are characterized by different degrees of demyelination and inflammation[24]. Immunohistochemistry/fluorescence for the proteins FCGR2B and HLA-DR in six MS lesion types, showed that Hs8 macrophages are scarce in non-active lesions and almost exclusively found in and in close proximity to blood vessels (Fig. 3H, I, J). Conversely, in active MS lesions, IBA1^pos^FCGR2B^pos^HLA-DR^pos^ cells (Hs8) were much more abundant in both blood vessels and the parenchyma (Fig. 3H, I). These data indicate that macrophages and/or monocytes with a CAM-like signature infiltrate the brain parenchyma during active lesion formation or that parenchymal microglia adopt a CAM-like signature.

Expression of heat-shock-protein (HSP) genes, IEGs and genes associated with stress response was enriched in cluster Hs6. Since these findings are in line with the enrichment of HSP-genes and IEGs in total tissue NAWM compared to CWM (Supplementary Fig. S1B), this strongly suggest that HSP gene and IEG expression of macrophages in NAWM compared to NACT is not an intrinsic property of WM macrophages, but rather is MS pathology associated.

Expression of microglia marker genes was enriched in the majority of the cells, but depleted in clusters Hs6, Hs7 and Hs8 (Fig. 3D). The expression of typical homeostatic microglia markers including *CX3CR1*, *P2RY12 and IRF8* was enriched in clusters Hs1, Hs2, Hs3 and Hs5, suggesting that these clusters represent homeostatic microglia (Fig. 3F, Supplementary Table S4). The expression of genes associated with the complement system was enriched in cluster Hs3 (Fig. 3F). This cluster was most abundant in NACT samples, but also slightly enriched in WML compared to NAWM (Fig. 3E, Supplementary Fig. S2C). The expression of pro-inflammatory genes was enriched in cluster Hs4, this cluster was equally abundant in all groups (Fig. 3E, Supplementary Fig. S2C). The expression of myelinogenic, developmental and disease-associated microglia genes (*FTL*, *SPP1*, *ASAH1* and *GPNMB*), that have activated/phagocytic microglia phenotypes in mice, was enriched in cluster Hs7 (Fig. 3F, G, Supplementary Table S4). These data indicate the cells in cluster Hs7 are microglia, associated with demyelination or other myelin-associated processes.

### Activated/phagocytic microglia arise during demyelination

To validate the hypothesis that Hs7 cells are associated with demyelination, we profiled brain macrophages in the cuprizone mouse model of de- and remyelination. Mice were fed with cuprizone or control diet for 3 or 5 weeks and macrophages were isolated for scRNAseq from the whole brains with exclusion of the olfactory bulb, cerebellum and brainstem (Fig. 4A). Cx3cr1^pos^ cells were sorted (Supplementary Fig S3A) from a Cx3cr1^lox-stop-tdTomato^ reporter mouse strain one month after tamoxifen administration to activate the reporter and were clustered based on their gene expression profiles (Fig. 4B, Supplementary Fig. S3A, B). Significant changes in cluster distribution between the groups were identified (Fig. 4C). Marker genes were identified by differential gene expression analysis between all clusters (Fig. 4B, Supplementary Table S6). Expression of genes specific for CAMs (*F13a1*, *Lyve1*, *Cd163*) and monocytes (*H2-Ab1*, *H2-Aa*, *Cd74*, *Lyz*) was enriched in clusters Mm7 and Mm8, respectively, and these clusters were equally abundant in all groups (Fig. 4C, Supplementary Fig. S3C, D). Expression of genes associated with microglia homeostasis, such as *P2ry12* and *Tmem119,* was enriched in clusters Mm1 and Mm2 and depleted in Mm3, Mm4, Mm7 and Mm8 clusters (Fig. 4D, Supplementary Fig. S3C, D, Supplementary Table S6). Two clusters with an increased abundance in the demyelination groups (D3, D5) were identified: Mm3 and Mm4 (Fig. 4C). Marker genes of Mm3 were *Lpl*, *Apoe*, *Spp1* and *Axl* (Fig. 4D, Supplementary Fig. S3C, D), genes previously identified in microglia in several disease models and during development and myelinogenesis[14, 18, 39]. Expression of genes associated with the interferon response, such as *Ifit3*, *Stat1* and *Irf7* was enriched in the Mm4 cluster (Fig. 4D, Supplementary Fig. S3C, D). A cluster of cells (Mm5) highly expressing *Phlda3* and *Cdkn1a* (Fig. 4D, Supplementary Fig. S3D), associated with apoptosis, was identified which was most abundant after 3 weeks demyelination (Fig. 4C). Additionally, a cluster of cells (Mm6) associated with proliferation was identified, as indicated by high expression of *Mki67 and Top2a* (Fig. 4D, Supplementary Fig. S3D), that also increased in abundance after 3 weeks demyelination. This finding was supported by immunofluorescent stainings for the proliferation marker Ki67 with IBA1, which revealed proliferating macrophages in the CP of demyelinated mouse brains after 3 weeks cuprizone (Fig. 4D). Remarkably, in the remyelination group (2 weeks after discontinuation of the cuprizone diet), the cluster distribution was very similar to the control condition (Fig. 4C), indicating that the Mm3 and Mm4 phenotypes may be reversible. The identification of the apoptotic (Mm5) and proliferation (Mm6) clusters in the demyelination groups suggests that Mm3 and Mm4 microglia might be (partly) replaced by proliferating microglia.

**Figure 4.**
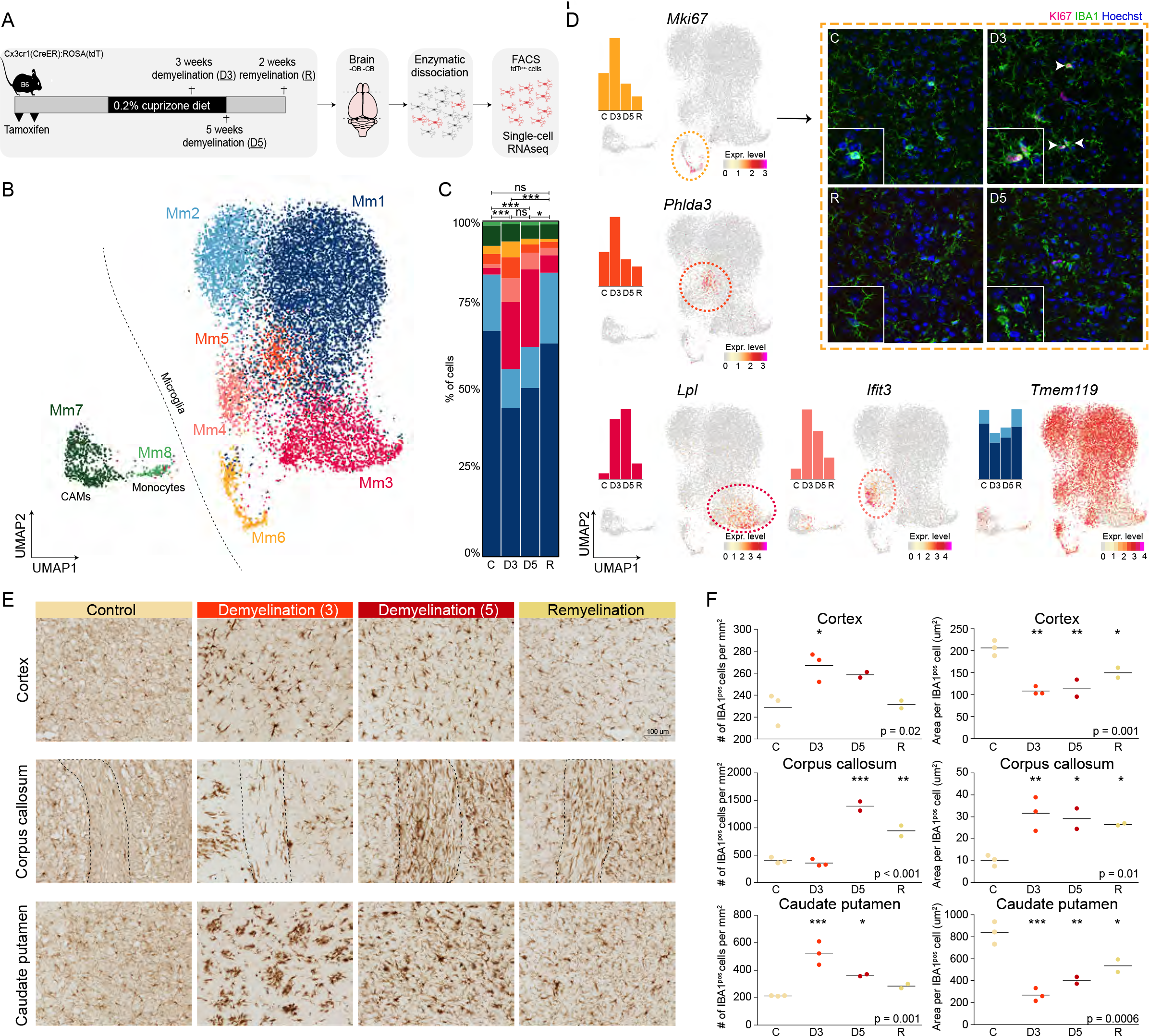
Activated/phagocytic microglia arise during demyelination. (**A**) Workflow of the experiment. (**B**) UMAP depicting 12,885 cells. Colors depict unsupervised clustering. (**C**) Bar plot depicting cluster distribution per group with statistical analysis. (**D**) UMAPs depicting gene expression levels of *Mki67*, *Phlda3*, *Lpl*, *Ifit3* and *Tmem119*. Immunofluorescent staining of caudate putamen region. Hoechst (blue; nuclei), IBA1 (green; macrophages), Mki67 (magenta; Mm6). Arrowheads indicate IBA1^pos^KI67^pos^ cells. (**E**) representative images of immunohistochemistry for IBA1 in the cortex, corpus callosum and caudate putamen. (**F**) Quantification of immunohistochemistry of 2-3 mice per group. *: p < 0.05; **: p < 0.01; ***: p < 0.001. Abbreviations: C = control; D3 = 3-weeks demyelination; D5 = 5-weeks demyelination; R = remyelination.

Changes in microglia proliferation and transcriptomic profiles were confirmed by altered macrophage morphology (Fig 4E, 4F right panel) and density (Fig 4E, 4F left panel) in situ. Immunohistochemistry for the macrophage marker IBA1 showed that macrophage morphology was extremely affected in the cortex and caudate putamen after 3 weeks on the cuprizone diet (D3) and nearly completely restored after 2 weeks remyelination. In the corpus callosum, macrophage morphology was changed the most after 5 weeks of cuprizone diet (D5), and this effect was still moderately present after 2 weeks remyelination, indicating that not all brain regions are affected at the same time, likely due to differences in myelin content (Fig. 4E, F). Taken together, these data indicate that the activated/phagocytic profile (Mm3) arises during demyelination and disappears after remyelination.

### Activated/phagocytic microglia are present in lesion types with ongoing demyelination

Next, we compared the macrophage subtypes identified in the cuprizone mouse study with the MS subtypes. Hs1, Hs2 and Hs3 macrophage clusters were significantly enriched for marker genes of the Mm1 and Mm2 clusters from the mouse study and depleted for Mm3, Mm7 and Mm8 marker genes (Fig. 5A). Mm7 (CAMs) and Mm8 (monocytes) marker genes were particularly enriched in Hs8 and Hs10 (neutrophils), again showing that the Hs8 cluster may contain CAMs or monocytes rather than microglia. Human cluster Hs6 was not significantly enriched in any of the mouse clusters, indicating that this is a specific feature of MS pathology that is not recapitulated in the healthy mouse brain or in response to cuprizone-induced demyelination. Conversely, the mouse apoptotic (Mm5) and proliferating (Mm6) clusters were not detected in the human data. Human cluster Hs7 was significantly enriched for expression of the Mm3 mouse cluster that both showed an activated/phagocytic profile (Fig. 5A), and cells in clusters Hs7 and Mm3 had comparable differential gene expression profiles compared to homeostatic cells, clusters Hs5 and Mm1 (Fig. 5B). Mm3 microglia appeared specifically in relation to demyelination in the cuprizone mouse model (Fig. 4C), therefore the association of Hs7 microglia with particular MS lesion types was investigated. Immunohistochemical and immunofluorescent stainings for two marker genes of cluster Hs7 in six different types of MS lesions in 3-11 independent donors showed that ASAH1^pos^ and LGALS1^pos^ cells were particularly observed in pre-active, active and mixes active/inactive lesions, confirming that Hs7 macrophages with an activated/phagocytic profile are associated with (active) demyelination (Fig. 5C, D, Supplementary Fig. S4).

**Figure 5:**
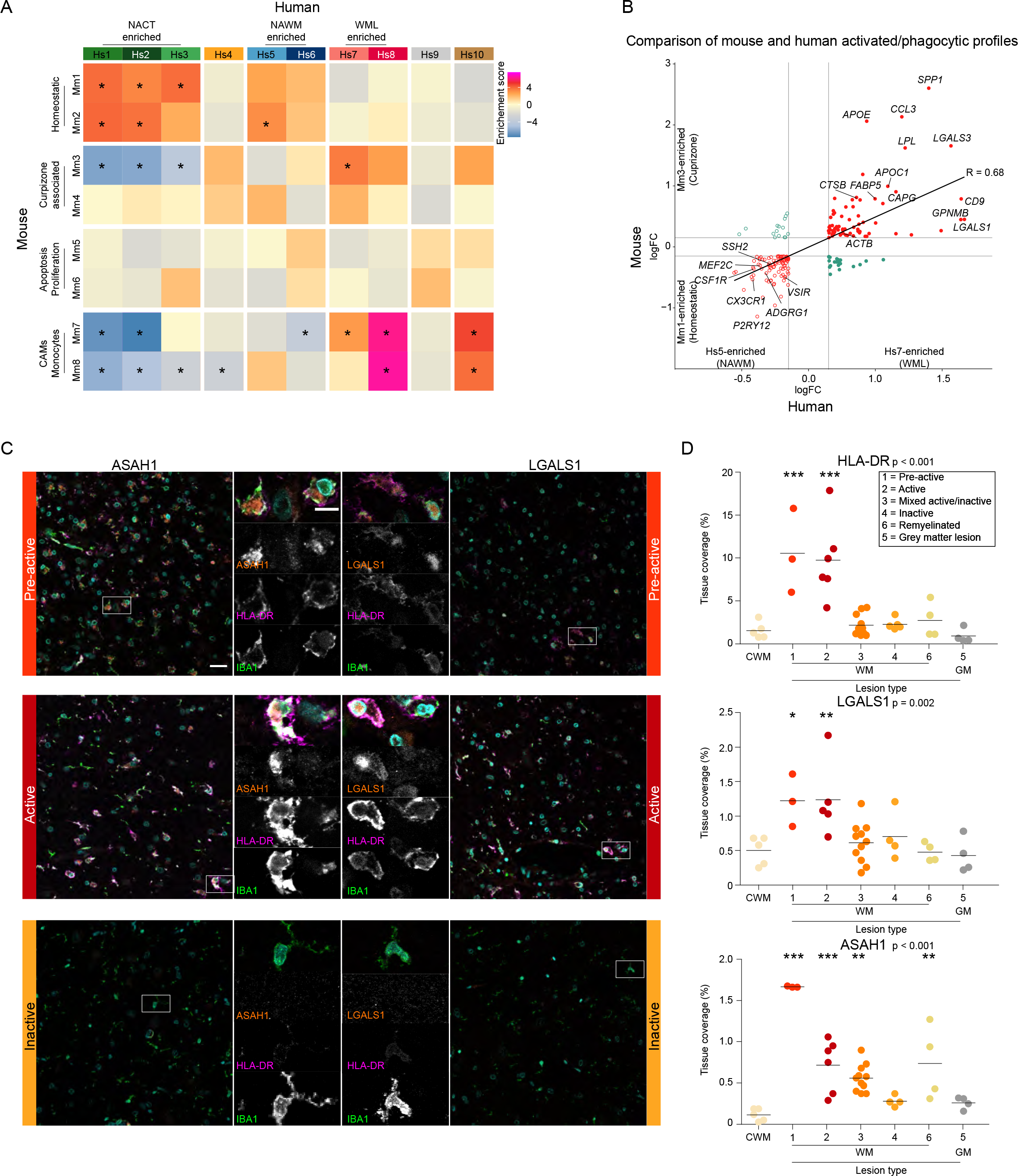
Activated/phagocytic microglia are present in lesion types with ongoing demyelination. (**A**) Heatmap depicting expression weighted gene set enrichment scores of cluster markers derived from the cuprizone experiment in the human MS dataset. (**B**) Fourway plot depicting gene signature of human Hs7 cluster compared to Hs5 cluster (x-axis) versus mouse Mm3 versus Mm1 cluster (y-axis). (**C**) Images of immunofluorescent staining for ASAH1 (orange, left; MS-4), LGALS1 (orange, right; Hs7), HLA-DR (magenta; Hs7), Hoechst (cyan; nuclei) and IBA1 (green; macrophages) in a pre-active, active and inactive lesion. Scale bar is 25 μm (overview), and 10 μm (zoom). (**D**) Quantification of immunohistochemistry for HLA-DR, LGALS1 and ASAH1 in CTR donors and MS donors amongst 6 different lesion types in 3-11 donors per group. *: p < 0.05; **: p <0.01; ***: p <0.001. Abbreviations: CWM = control white matter (CTR donor); NAWM = normal appearing white matter (MS donor); WML = white matter lesion (MS donor); NACT = normal appearing cortical tissue (MS donor); CAM = CNS-associated macrophages.

Taken together, this study indicates the presence of an early stress response in macrophages in NAWM of MS brains, and that the activated/phagocytic microglia profile in WML is an early but persistent response to disease, associated with ongoing demyelination during lesion formation and progression.

## Discussion

Here, transcriptomic profiling of white matter from control donors, and NAWM and WMLs from MS brain tissues is performed to detect early changes that may underlie MS disease onset. This total tissue transcriptomic profiling pointed towards changes in brain macrophages in NAWM tissue. Next, we performed single-cell RNAseq of brain macrophages to investigate how these cells are affected in NAWM and WML tissue. For this, fresh post-mortem brain tissue was obtained within 12 hours after autopsy. Hence, it was not possible to process multiple donors at the same time, possibly leading to batch effects induced by technical artefacts or post-mortem variation. Therefore, we processed NAWM, WML and NACT from the same donors (in one batch), to overcome inter-donor variation that may be confounded with technical variation or other (post-mortem) artefacts. Rather than including brain tissue from control donors, NACT was included as a reference, non-diseased brain region to obtain sufficient heterogeneity in the single-cell RNAseq data to identify the transcriptomic profiles of NAWM- and WML-specific macrophage subpopulations. Additionally, MS donors die relatively young (often 55-70 years of age), whereas “healthy” control (CTR) donors generally die due to aging (> 80 years of age), hampering an age-matched design. Thus, using brain tissue from CTR donors would create an aging bias in the dataset, and it would take years to obtain sufficient numbers of fresh young CTR brain tissues which may make the data even more prone to technical variation. We acknowledge that macrophage profiles differ in grey and white matter tissue, which we also show in our dataset. Differences in microglia transcriptomes from grey and white matter was also reported by van der Poel et al. (2019) in both control and MS brains[30]. Nevertheless, we saw a large spectrum of macrophage/microglia heterogeneity in human MS brains, and besides homeostatic microglia, we observed signatures that were nearly exclusively observed in NAWM and WML and displayed disease-associated signatures that partly overlap with those found in disease models, during development and in other neurodegenerative diseases[11, 14, 18, 33, 38].

Using scRNAseq, a cluster of macrophages was identified that was particularly abundant in NAWM samples and showed enriched expression of heat shock proteins and immediate-early genes (Hs6). These genes have been reported to be induced by ex vivo activation of macrophages, such as incubation steps at 37 °C[27]. Importantly, our cell isolation was performed at 4° C throughout the entire procedure. This stress-related gene signature was also identified in the total tissue bulk RNAseq dataset, both by us and partly by Melief et al. (2019)[28], suggesting that these genes are not induced by cell isolation procedures but are an early feature of MS pathology that already occurs before the onset of demyelination. Even more, these stress-associated macrophages did not appear in cuprizone-induced demyelinated mice, confirming that the stress macrophage cluster does not arise in response to demyelination itself but arises in MS brains before demyelination occurs. These data show that brain macrophages display an early stress response prior to demyelination, suggesting that these are amongst the first cells to be involved in the onset of demyelination. Alternatively, the stress response in brain macrophages may represent a protective response of these cells, aiming to prevent lesion formation in that tissue area.

Two clusters of macrophages were identified that were particularly enriched in demyelinated WML samples. Cluster Hs8 showed enriched expression of several well-known CAM genes, such as *F13A1* and *LYVE1*[16]. Additionally, cluster Hs8 was significantly enriched for marker gene expression of the CAMs and monocytes that were identified in the cuprizone dataset. The role of microglia and infiltrating macrophages and monocytes in MS brains is complicated by the fact that they are remarkably similar in case of disease[4, 13]. While under homeostatic conditions microglia express certain unique markers, including TMEM119, these microglia markers are downregulated in disease[4, 13]. At the same time, following exposure to the CNS microenvironment of microglia-deficient mice, CNS-infiltrating macrophages adopt a microglia-like phenotype[3]. Therefore, despite the detection of marker genes for CAMs, from these data we cannot definitively conclude whether Hs8 macrophages indeed represent CAMs or that they are microglia that express CAM marker genes in response to pathology. In mice, it was shown that microglia outnumber CAMs and dominate the CNS lesion in response to demyelination[29], making it more likely that these Hs8 macrophages originate from microglia rather than CAMs. On the other hand, blood-brain-barrier (BBB) dysfunction is a well-known feature present in WMLs. BBB dysfunction makes the CNS prone to increased infiltration of macrophages and/or monocytes[7]. In non-active lesions and control brain tissue, Hs8 macrophages were rare and almost exclusively observed near blood vessels. In (pre)-active and chronic lesions, Hs8 macrophages were more frequent and also present in the parenchyma, suggesting that Hs8 macrophages may come from blood vessels and invade the lesion parenchyma, or, alternatively, that resident microglia switch to an Hs8 phenotype.

Cluster Hs7 showed an activated/phagocytic microglia signature that was similar to profiles reported in mouse models of disease[16, 18], human AD[11] and during fetal/neonatal development and myelinogenesis[14, 19, 38]. Hs7 microglia were particularly detected in lesion types with ongoing active demyelination. A similar gene expression profile was identified in the cuprizone dataset (Mm3), confirming their association with active demyelination. Additionally, these Mm3 microglia were no longer abundant when the brain was fully remyelinated. During mouse development, a similar subset of microglia (axon tract associated microglia) appears in regions that become heavily myelinated and disappears when myelination is complete[14]. Another study also showed that a similar microglia subtype (CD11C^pos^) is abundant in areas of primary myelination and is a critical source of IGF1-driven myelination[38]. These data indicate that in WMLs of MS brains, a phagocytic/activated microglia profile (Hs7) emerges that is similar to those found in the developing brain and is associated with myelin-processes.

Taken together, our data reveals that macrophages in MS brains adopt a diverse range of phenotypes. Furthermore, significant changes were observed in NAWM tissue, indicating that prior to apparent lesion formation (demyelination), brain macrophages are already responding to environmental cues. These data offer insight into early disease-associated changes in MS brain tissue in relation to lesion development and progression, that may provide therapeutical targets to prevent or reduce demyelination.

## Materials and methods

## Bulk RNAseq experiment

### Post-mortem frozen human brain tissue and lesion classification

Snap frozen white matter (WM) MS brain tissue was obtained from the Netherlands Brain Bank (NBB, Amsterdam, The Netherlands) (n = 117). From these MS donors the diagnosis progressive MS was confirmed. Age and gender matched post-mortem brain tissues from control donors were obtained from the Edinburgh Brain and Tissue Bank (EBTB, Centre for Clinical Brain Sciences, University of Edinburgh) (n = 35). WM tissue blocks from MS donors and controls were characterized by immunohistochemistry. Per tissue block every first and last two cryosections (5µm) were used for immunohistochemical stainings, while the intermediate part was used for bulk mRNA sequencing. For sample characterization, the presence or absence of lesions or inflammation was confirmed by scoring demyelination and immune activation through the markers proteolipid protein (PLP1) and Human Leukocyte Antigen – DR isotype (HLA-DR) respectively. An HLA-DR and PLP1 score ranging from 0 to 10 was assigned as previously described [20, 36] (Supplementary Fig. S1A). Tissues with PLP1 score 0 (not intact myelin) and HLA-DR score higher than 5 (high macrophage activation) were included in the study as WM lesions, while all tissues with PLP1 score 10 (intact myelin) and HLA-DR score lower than 4 (low macrophage activation) were included in the study as NAWM. In addition, the PLP1 staining was used to determine if the white matter samples were contaminated with grey matter areas. Samples with grey matter contamination were excluded for further analysis.

Cryosections were fixed with acetone (for HLA-DR) or 4% paraformaldehyde (for PLP1) for 10 minutes and 70% ethanol for 5 minutes, followed by a 3 minutes incubation in PBS. Next, endogenous peroxidase activity was suppressed using 0.3% H_2_O_2_, followed by blocking in 5% normal horse serum (NHS) in PBS for 30 minutes. Sections were incubated overnight with mouse anti-human HLA-DR (eBioscience, 17-9956-42, 1:750) or mouse anti-human PLP1 (Serotec, MCA839G, 1:500), followed by incubation with biotin-conjugated horse-anti mouse secondary antibody (Vector, BA-2000-1.5, 1:400) for 2 hours at room temperature. After a 30 min incubation with the avaidin-biotin solution (ABC, Vectastain ABC kit, Vector, PK-6100), the complex was visualized with DAB in PBS containing 0.03% H_2_O_2_. Subsequently, haematoxylin was used as nuclear counterstain, followed by mounting in DePeX (Serva, 18243). Images were digitalized using a NanoZoomer 2.0-HT Digital slide scanner C9600 (Hamamatsu Photonics). Sections were scored for HLA-DR and PLP1 using an Axioskop microscope (Carl Zeiss).

### Bulk RNAseq of frozen brain tissue

WM snap frozen tissue blocks were classified as WML containing either an active, mixed active/inactive or remyelinated lesion (for lesion classifications of each sample based on Luchetti et al. 2018[24] see Supplementary table S1), NAWM and CWM (Supplementary Fig. S1A). Tissue sections immediately surrounded by the sections used for classification was collected for RNA extraction and whole tissue 3**’** mRNAseq. This concerned 20 10µm cryosections that were collected in 1 mL QIAzol (Qiagen, 79306) and lysed and homogenized with a syringe. RNA was isolated using the RNeasy Lipid Tissue mini kit (Qiagen, 74804) following manufacturer’s instructions. RNA was eluted in 40 µl RNase free water, RNA concentrations and integrity were measured on a Bioanalyzer 2100 (Aligent). Samples with a RIN value > 4 were included for downstream analysis. cDNA libraries were obtained with the QuantSeq 3’ mRNA-Seq Library Prep Kit FWD for Illumina (Lexogen, 01596) according to the manufacturer’s protocol. Samples were sequenced on an Illumina NextSeq 500 Sequencing System with NextSeq 500/550 High Output Kit v2.5 (Illumina, 20024906).

### Bulk RNAseq analysis

For bulk analysis, raw reads were aligned to the GRCh37 reference genome from Ensemble with Hisat (v0.1.5). Aligned reads were sorted with samtools (v1.2) and counted with HTSeq (v0.6.1). FastQC (v0.11.3) and Picard (v1.130) were used to perform quality control. Raw count matrices were loaded in R and annotated by converting the ensemble IDs to gene symbols using the corresponding .gtf file. Lowly expressed genes were filtered using a data-adaptive flag method for RNA-sequencing (DAFS[10]). Only genes with > 1 counts in at least 2 samples were included in the analysis. To determine whether donors from the same group would cluster together, the count matrix was normalized with the blinded variance-stabilizing method from DESeq2 from Bioconductor and mitochondrial genes were removed prior to this analysis. A negative binomial generalized log-linear model was used to model gene expression levels and differentially expressed genes were determined using a likelihood ratio test [31]. Thresholds were set at abs(logFC) > 1 and adjusted-p < 0.05. Principal component analysis was performed on VST-transformed counts. Visualizations were made with the CRAN package ‘ggplot2’. For WGCNA analysis, VST-transformed counts obtained from DESeq2 were used as input[21, 23]. Signed WGCNA was performed using biweight mid-correlations and the maximum number of excluded outliers was restricted to 10%[22]. Gene ontology analysis was performed with MetaScape[41]. Cell (sub)type enrichment analyses were performed with expression weighted cell type enrichment analysis (EWCE)[34] using CTR donors from the snRNAseq dataset from[11] as a reference. For DEG gene-sets, logFC was used for gene ranking in EWCE, for weighted gene co-expression network analysis (WGCNA) the module member ship scores.

## Human brain macrophage experiment

### Fresh post-mortem human MS brain tissue

Of 5 donors, fresh post-mortem brain tissue was obtained from the Netherlands Brain Bank (NBB). Immediately after autopsy, the tissue was transported from Amsterdam to Groningen in HBSS with phenol red (Thermofisher Scientific, 14170-088) supplemented with 15 mM HEPES (Lonza via Westburg, LOBE17-737E) and 0.6% glucose (Sigma Aldrich, G8769). Of each donor three samples were obtained: i.e. 1) normal-appearing cortical tissue mainly consisting of a mixture of grey- and white matter (NACT); 2) normal-appearing white-matter (NAWM); 3) white-matter lesion (WML) tissue with the surrounding perilesional WM area (Fig. 3A). Lesions were selected by the NBB based on post-mortem magnetic resonance imaging (MRI) together with detailed macroscopic observations. Due to limited availability of fresh post-mortem MS tissues and the relatively small size of the tissue samples with MS lesions, all obtained tissue was used for single cell sequencing and, therefore, the MS lesion type and HLA-DR activity within the NAWM are unknown.

Furthermore, due to limited control donors of similar ages, tissue from aged matched control donors was not available. To correct for age as a batch effect and to allow for within donor comparisons and generate sufficient contrasting signatures to identify disease-associated signatures, we included multiple samples from each donor. Besides WML and NAWM tissue, normal appearing cortical tissue (NACT) was used as an internal reference group, since pathology in GM is known to be distinct from WM pathology. Informed consent to perform autopsies and the use of tissue and clinical data for research purpose were obtained from donors and approved by the Ethical Committee of the VU University Medical Center (VUmc, Amsterdam, The Netherlands).

### Macrophage isolation from human brain tissue

Human macrophages were isolated as described previously[9]. In brief, meninges were removed and the tissue was mechanically dissociated using a glass tissue homogenizer. A cell suspension was obtained via filtering through 300-µm and 106-µm sieves. Myelin was removed by a 24% Percoll gradient (Fisher Scientific, 17-0891-01) in 10x HBSS (Gibco, 14180-046) and phosphate buffered saline (PBS) density gradient centrifugation and followed by a second centrifugation step containing a 60% and 30% Percoll layer with PBS on top. The interphase between the Percoll layers was collected and contained the immune cells. Fc receptors were blocked with human Fc receptor-binding inhibitor (eBioscience, 14-9161-73). For FACS, cells were incubated for 20 min with anti-human CD11B-PE (BioLegend, 301306) and anti-human CD45-FITC (BioLegend, 304006) and washed with HBSS without phenol red (Thermofisher Scientific, 14175-053). The cells were passed through a 35-μm nylon mesh, collected in round bottom tubes (Corning 352235), stained with DAPI (Biolegend, 422801) and DR (Thermofisher Scientific, 62251) and sorted using a Beckman Coulter MoFloAstrios, Beckman Coulter MoFloXDP or Sony SH800S cell sorter. Human macrophages were sorted as DAPI^neg^DR^pos^CD11B^pos^CD45^pos^.

### Single-cell RNA sequencing mouse and human cells

The single cell cDNA libraries were constructed using the Chromium Single Cell 3’ Reagents Kit v2 and corresponding user guide (10x Genomics) and sequenced on a NextSeq 500 at the sequencing facility in the UMCG up to a depth of ∼20,000 reads/cell.

### scRNAseq data analysis

Raw reads were aligned to the GRCh38 or GRCm38 genome for human and mouse samples, respectively, using Cell ranger (v3.0.0) with default settings. Raw count files were loaded into R (v3.6) and barcode filtering was performed with thresholds at >600 unique molecular identifiers (UMIs) for mouse cells and >400 for human cells[26]. The multiplet rate mentioned in the 10x Genomics User Guide was used to set an upper threshold per sample for the number of UMIs per cell. Cells with a mitochondrial content >5% were removed from the dataset.

For the mouse dataset, the count files from different conditions were merged into one and further analyzed with Seurat[35]. The data were normalized by dividing the counts of each gene by the total sum of counts per cell and multiplied by a scale factor of 10,000 and log-transformed. Highly variable genes (HVGs) were calculated using the ‘VST’ method with default settings. The data was scaled and heterogeneity associated with number of UMIs and mitochondrial and ribosomal content were regressed out, then the data was clustered. Differential gene expression analysis was performed with MAST[8].

For the human dataset, count matrices of the three brain regions per donor were merged into one file per donor and normalized using the same method as for the mouse data. HVGs were determined using the VST method. The datasets from the donors were integrated using canonical correlation analysis[35]. The data was scaled and heterogeneity associated with number of UMIs, mitochondrial content, sex and ribosomal content were regressed out. The data was clustered using the graph-based clustering method implemented in Seurat with default settings. Differential gene expression was performed on the unintegrated data using logistic regression with donor as a latent variable. Geneset module scoring was performed using the AddModuleScore function in Seurat. Cell (sub)type enrichment analyses were performed with EWCE using the human scRNAseq as a reference dataset and marker genes of the cuprizone scRNAseq clusters ranked by logFC[34]. Statistical analysis of cluster distribution changes between groups was performed using chi-squared tests in R.

### Gene sets from literature

From [33], EV7 was downloaded and genes with a p_val_adj < 0.05 and abs(logFC) > 0.15 were selected. From [14], table S1 was downloaded and marker genes from cluster 4 were selected. From [18], table S2 was downloaded and upregulated genes of “Microglia3” with a p_val_adj < 0.05 were selected. From [38], dataset EV1 was downloaded and DEGs between neonatal CD11c^pos^ microglia and neonatal microglia with abs(logFC) > 2.5 and FDR < 0.05 were used. From [11], DEGs between AD1 microglia and homeostatic microglia (abs(logFC) > 0.15 and p_val_adj < 0.05) were used. From [19] cluster markers of cluster 8 were used (abs(avg_logFC) > 0.25 and p_val_adj < 0.05).

## Paraffin-embedded human brain tissue

Formalin-fixed paraffin-embedded, well-characterized tissues from 5 controls and 14 MS donors were obtained from the NBB. Lesions were classified based on HLA-DR and PLP1 immunohistochemistry according to the system for lesion classification from the NBB[24], resulting in the following lesion types indicated by a number: pre-active (1), active (2), mixed active/inactive (chronic) (3), inactive (4) and shadow plaques or remyelinated lesions localized in WM (6) and GM lesions (5). Paraffin blocks were cut into 6-µm thick sections. After deparaffinization and heat-induced antigen retrieval in 10 mM sodium citrate (pH 6.0) with 0.05% Tween for 10 minutes in a microwave, sections were divided in two series. We used one series of sections for immunohistochemistry and subsequent quantification and MS lesion scoring, using the Avidin-Biotin Complex (ABC) method followed by a DAB staining. The other series of sections we used for triple immunofluorescence stainings with various antibody combinations.

### Immunohistochemistry

After deparaffinization and antigen retrieval, endogenous peroxidase was blocked using H_2_O_2_/PBS 0.3% for 30 minutes. After washing with PBS, tissue sections were incubated for 30 minutes with 2% normal serum (NS) and 2% bovine serum albumin (BSA). After a short wash with PBS, sections were incubated overnight at 4°C with the primary antibody in PBS containing 1% NS and 1% BSA (Table 1). The following primary antibodies were used (Table 1): anti-PLP1 (AbD Serotec, MCA839G, dilution 1:100), anti-HLA-DR (eBioscience, 14-9956, dilution 1:500), anti-FCGR2B (Bioorbyt, orb44658, dilution 1:350), anti-ASAH1 (Abcam, ab74469, dilution 1:500), anti-LGALS1[2, 17] (dilution 1:100). Next day, sections were washed with PBS and incubated for 2 hours with the appropriate biotinylated secondary antibodies (anti mouse or rabbit, Vector Labs, dilution 1:400; Table 1), followed by a 30 minutes incubation with the ABC solution (Vectastain elite kit, PK-6100). After rinsing with PBS, the sections were incubated for 10 minutes in DAB and 0.03% H_2_O_2_. Finally, the sections were counterstained with cresyl violet, dehydrated, and mounted with DepeX (Serva, 18243). Images were acquired using a NanoZoomer 2.0-HT Digital slide scanner C9600 (Hamamatsu Photonics). Images of the HLA-DR and PLP1 stainings were used for scoring the MS lesion, the results of the FCGR2B, ASAH1 and LGALS1 stainings were quantified as follows. From every type of lesion five 40x magnified images were extracted per donor. To measure FCGR2B activity near blood vessels, 80X magnified images were extracted from the scans. Analysis was performed using FIJI by first applying colour deconvolution using the H & DAB method. From the DAB channel a binary image was made using manual thresholding and the area fraction was measured in ImageJ. Statistics were performed with a two-way ANOVA for location and lesion type, followed by Tukey post-hoc test for lesion types (Fig. 3I, J). Statistics on DAB total tissue coverage quantifications were performed with a one-way ANOVA followed by Dunnet’s test for comparison of each group versus the CWM group (Fig. 5D).

**Table 1.**

| Immunohistochemistry |  |  |  |
| --- | --- | --- | --- |
|  | Company | Cat # | Dilution |
| **Antibody (primary)** |  |  |  |
| PLP1 (clone plpc1) | AbD Serotec | MCA839G | 1:500 |
| HLA-DR (clone LN3) | eBioscience | 14-9956 | 1:750 |
| FCGR2B (datasheet CD32B) | Biorbyt | orb44658 | 1:350 |
| ASAH1 | Abcam | ab74469 | 1:500 |
| LGALS1 |  |  | 1:100 |
| Claudin5 (clone 4C3C2) | Thermo Fisher Scientific | 35-2500 | 1:250 |
| IBA1 | Wako | 019-19741 | 1:1000 |
| IBA1 | Abcam | ab5076 | 1:750 |
| Ki67 (Clone B56) | BD Pharmingen | 556003 | 1:400 |
| **Antibody (secondary)** |  |  |  |
| Biotin-Donkey anti-Rabbit IgG | JacksonImmunoRese<br>arch | 711-065-152 | 1:300 |
| Cy3 Goat anti-Mouse IgG | JacksonImmunoRese<br>arch | 115-165-003 | 1:200 |
| Donkey anti-Mouse, Alexa<br>Fluor™ 594 | Thermo Fisher<br>Scientific | A21203 | 1:300 |
| Donkey anti-Goat, Alexa Fluor™<br>633 | Thermo Fisher<br>Scientific | A21082 | 1:300 |
| Streptavidin, Alexa Fluor™ 488 | Thermo Fisher<br>Scientific | S11223 | 1:300 |
| **FACS** |  |  |  |
| **Antibody (primary)** | **Company** | **Cat #** | **Dilution** |
| FITC anti-human CD45 | BioLegend | 304006 | 1:25 |
| PE anti-human CD11b | BioLegend | 301306 | 1:40 |

### Immunofluorescence

After deparaffinization and antigen retrieval as described above, sections were incubated for 10 minutes in a Sudan Black solution (0.3% Sudan Black/Ethanol 70%) to quench autofluorescence followed by dipping the slides 3 times in ethanol 70% to remove excess of Sudan Black. After washing with PBS, tissue sections were incubated for 30 minutes with 2% NS and 2% BSA. After a short wash with PBS, sections were incubated overnight at 4 °C with primary antibodies in PBS containing 1% NS and 1% BSA (Table 1). A mixture of goat anti-Iba1 (Abcam, ab5076, dilution 1:500) and mouse anti-HLA-DR (eBioscience, 14-9956, dilution 1:500) or mouse anti-CLDN5 (Thermo Fisher Scientific, 35-2500, 1:250) was in various separate experiments supplemented with antibodies raised in rabbit, namely anti-FCGR2B (Bioorbyt, orb44658, dilution 1:350), anti-ASAH1 (Abcam, ab74469, dilution 1:500), anti-LGALS1[2, 17] (dilution 1:100). After an overnight incubation at 4°C, the sections were rinsed and incubated with a biotinylated anti-rabbit antibody (anti-rabbit, Vector Labs, dilution 1:400), followed by an incubation of 1 hour at room temperature with donkey anti-mouse IgG (H+L) AlexaFluor™ 594 (Thermo Fisher Scientific A21203), donkey anti-goat IgG (H+L) AlexaFluor™ 633 (Thermo Fisher Scientific A21082) and Streptavidin, AlexaFluor™ 488 conjugate (Thermo Fisher Scientific S11223) all diluted 1:300 in PBS plus Hoechst stain (Sigma 14530, 5μM final concentration). Sections were washed with PBS and demi water, mounted with Mowiol and imaged on a Leica SP8X confocal laser scanning microscope (Leica Microsystems, Amsterdam).

## Animal experiment

### Cuprizone mouse model

C57BL/6J-Cx3cr1^tm2.1(cre/ERT2)Litt^Gt(ROSA)26Sor^tm14(CAG-tdTomato)Hze^ mice were used in a model for cuprizone-induced demyelination. The mice were bred in-house on a C57BL/6J background. In order to activate Cre-recombinase and express the tomato reporter in CX3CR1 expressing cells, 6-weeks-old animals received twice 500 mg/kg body weight tamoxifen (Sigma Aldrich, T5648-5G) dissolved in corn oil (Sigma Aldrich, C8267-500ML) via oral gavage with a 3-day interval. All animal procedures were approved by the local central authority for scientific procedures on animals (CCD) and performed in accordance to ethical regulations (AVD105002015360). Demyelination was induced in 8-week-old male mice via a diet containing 0.2% w/w cuprizone (Sigma Aldrich, C9012-25G). The chow diet was freshly prepared every week by mixing cuprizone with standard powder food and water, and stored at -20 °C. Animals were provided with this home-made chow three times a week *ad libitum*. Control animals received similarly prepared chow, but without cuprizone. The experimental timepoints were: early demyelination (3 weeks cuprizone diet), complete demyelination, start remyelination (5 weeks cuprizone diet) and remyelination following withdrawal of the cuprizone diet for 2 weeks. To reduce biological variation and limit the effect of individual animals on the data, macrophages from n=5 animals per group were pooled into one sample for scRNAseq. Additionally, tissues from 3 animals were collected for immunohistochemistry.

### Macrophage isolation from mouse brain tissue

Mice were sacrificed under deep anaesthesia (4% isoflurane with 7.5% O_2_) and perfused with cold PBS. After perfusion, brains were removed from the skull and kept in cold medium A (HBSS (Gibco, 14170-088) with 0.6% glucose (Sigma, G8769) and 7.5 mM HEPES (Lonza, BE17-737E)). Macrophages were isolated enzymatically from the whole brain minus olfactory bulb and cerebellum. Brains were minced on a glass slide into small pieces with a knife and transferred to a tube containing enzyme solution with: 2 mL PBS, 20 mg Protease from *Bacillus licheniformis* (Sigma Aldrich, P5380-1G) and 20 µL L-cysteine, incubated on ice for 15 minutes while mixing every 5 minutes. After enzymatic dissociation, the cell suspension was passed through a 100µm cell strainer (Corning, 21008-950), filled up with 15 mL enriched HBSS. Cells were pelleted by centrifugation for 10 minutes, 300 RCF at 4 °C. Myelin was removed by 24% Percoll-(Fisher Scientific, 17-0891-and PBS density gradient centrifugation for 20 minutes, 950 RCF at 4 °C. The cells were passed through a 35-μm nylon mesh, collected in round bottom tubes (Corning, 352235) and sorted using a Beckman Coulter MoFloAstrios cell sorter. DAPI^neg^tdTomatoRed^pos^ cells were collected for scRNAseq.

## Immunohistochemistry and immunofluorescence on mouse brain tissue

4% PFA-fixed frozen mouse brains were sectioned at a thickness of 16 µm at bregma -0.8. This region was selected because most cuprizone induced demyelination is expected between bregma -0.6 and -1 based on previous experience. Sections were stained using immunohistochemistry or immunofluorescence method. For both methods, heat-induced antigen retrieval with sodium citrate (pH 6) was applied to unmask the epitopes.

### Immunohistochemistry

After heat induced antigen retrieval as described above, sections were washed with PBS and incubated with 0.3% H_2_O_2_/PBS to block endogenous peroxidases for 30 minutes. Fc receptor blocking was performed for 1 hour using 5% normal goat serum dissolved in PBS with 0.3% Triton X-100. After washing with PBS, sections were incubated overnight at 4 °C with the primary antibody: IBA1 (Wako, 019-19741, dilution 1:1000) in PBS containing 1% normal goat serum and 0.3% Triton X-100. The next day sections were washed with PBS and incubated with a biotinylated secondary antibody: goat-anti-rabbit (Vector, BA1000, dilution 1:400), followed by a 30 minutes incubation with the ABC solution (Vectastain elite kit, PK-6100). After rinsing with PBS, the sections were incubated for 10 minutes in DAB and 0.03% H_2_O_2_. Finally, the sections were dehydrated, and mounted with DepeX (Serva, 18243). Images were acquired using a NanoZoomer 2.0-HT Digital slide scanner C9600 (Hamamatsu Photonics). Per sample 10X zoomed images were saved from the striatum (1 image), corpus callosum (1 image) and cortex (2 images). For quantification, areas were used and averaged per animal with sizes of twice 500 x 500 µm (striatum), 200 x 625 µm (corpus callosum) and four times 400 x 400 µm (cortex) respectively. IBA1^pos^ cells within these drawn areas were quantified using the FIJI plugin ‘cell counter’. Additionally, images were converted to grayscale (8-bit images) and the percentage of IBA1 positive pixels was measured by FIJI. A constant threshold of 160 was used for the quantification. Statistics were performed with a one-way ANOVA followed by Dunnet’s test for comparison of each group versus the control group.

### Immunofluorescence

After heat-induced antigen retrieval as described above, blocking was performed for 1 hour using 5% normal goat and normal horse serum. Sections were incubated overnight with the primary antibodies: Ki67 (BD Pharmingen, 556003, dilution 1:400) and Iba1 (Wako, 019-19741, dilution 1:1000) in order to visualize proliferating microglia. After incubation and washing with PBS tissues were incubated with a secondary antibody mix containing donkey anti-rabbit AF488 (Thermofisher Scientific, A21206, dilution 1:400) and goat anti-mouse IgG Cy™3 (Jackson ImmunoResearch, 115-165-003, dilution 1:200) for 1.5 hours. Additionally, nuclei were visualized by Hoechst. Images were acquired with a Leica SP8X confocal laser scanning microscope (Leica Microsystems, Amsterdam) using a HC PL APO CS2 40x/1.30 oil objective in a sequential order with optimized emission using a white light laser and excitation detection using gated HyD detectors.

## Data availability statement

The datasets supporting the conclusions of this article are available through Gene Expression Omnibus at https://www.ncbi.nlm.nih.gov/geo with accession number GSE179427.

## Acknowledgements

The authors would like to thank Geert Mesander, Johan Teunis and Theo Bijma from the UMCG Flow Cytometry Unit. For 10X Genomics library preparation, we appreciate the help and flexibility of the EXPIRE group (UMCG). Sequencing was performed with the excellent assistance from Diana Spierings and the ERIBA Research Sequencing Facility. We thank the Netherlands Brain Bank for sample collection and Corien Grit, Eline Sportel and Marissa Dubbelaar for preliminary analyses.

## Author contributions

BJLE and SMK conceived the study. NB, EN, RPB, SA and SMK performed the bulk RNAseq experiment. AM, EG, NB, QJ, LK, EW, MHCW and SMK isolated cells from fresh human brain samples. EG, LK and SMK operated the FACS. AM and ATEP performed the mouse experiment. EW genotyped the mice. MM performed microscopy. HJG provided antibody for IHC. AM and EG performed scRNAseq. EG performed bioinformatic analyses and visualized the data. AM and NB performed validation experiments. AM, EG, NB, BJLE and SMK interpreted the data. AM and EG wrote the manuscript with supervision of BJLE and SMK. All authors critically read and approved the manuscript.

---

**Supplementary Figure S1.**
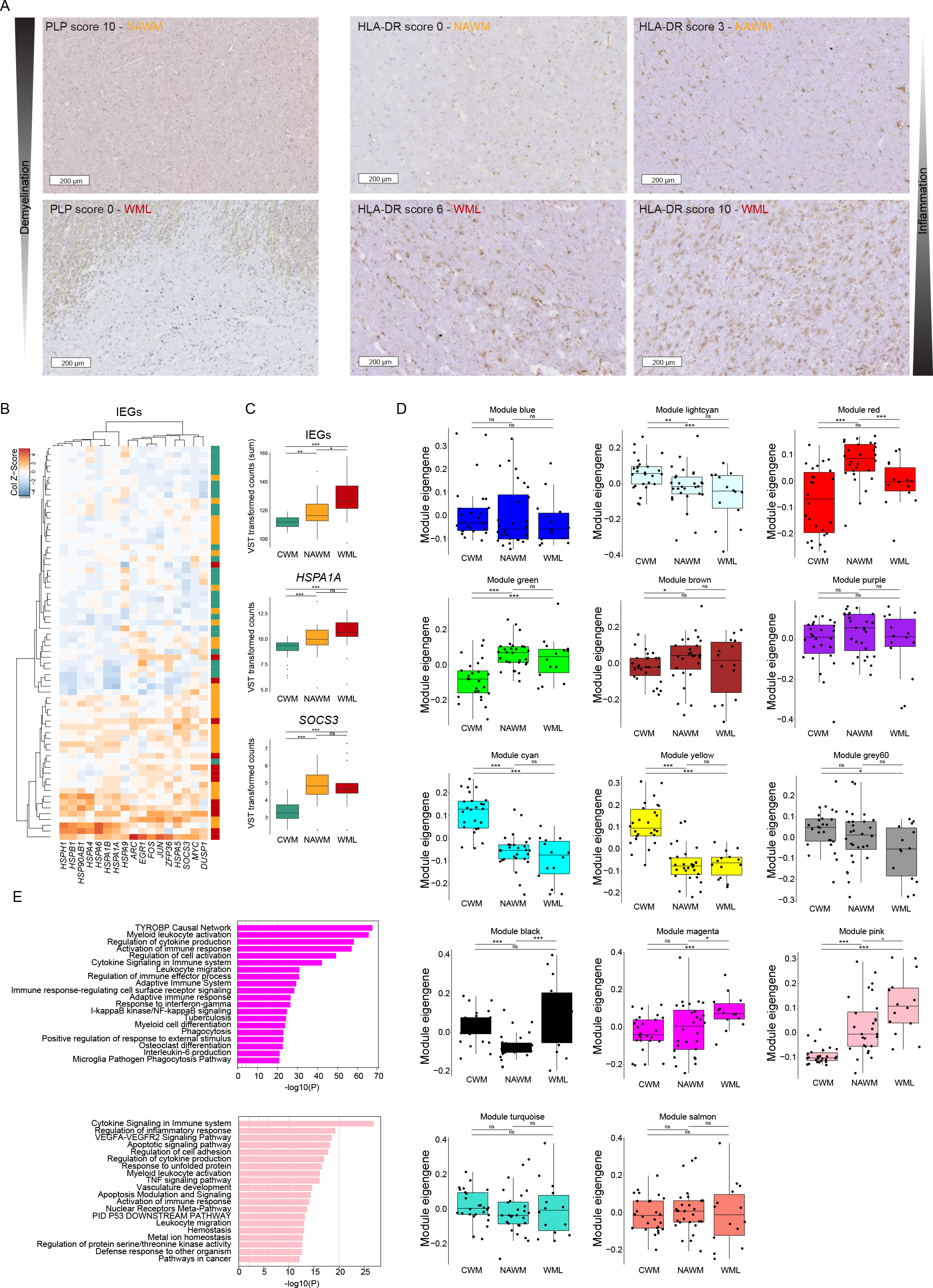
Supplementary data related to figures 1 and 2. (**A**) Representative images of DAB staining for PLP and HLA-DR that was used for sample scoring. (**B**) Heatmap depicting gene expression of a manually selected set of immediate-early genes. Rows and columns are ordered by hierarchical clustering. (**C**) Box plots depicting sum of IEG expression (top), *HSPA1A* expression level (middle) and *SOCS3* expression level (bottom). (**D**) Box plots depicting module eigengenes per sample group derived from the WGCNA of 68 white matter brain tissue samples. **(E**) Bar plots depicting gene ontology analysis of pink and magenta modules. *: p <0.01; **: p < 0.001; ***: p < 0.0001. Abbreviations: CWM = control white matter (CTR donors); NAWM = normal appearing white matter (MS donors); WML = white matter lesion (MS donors); IEGs = immediate early genes.

**Supplementary Figure S2.**
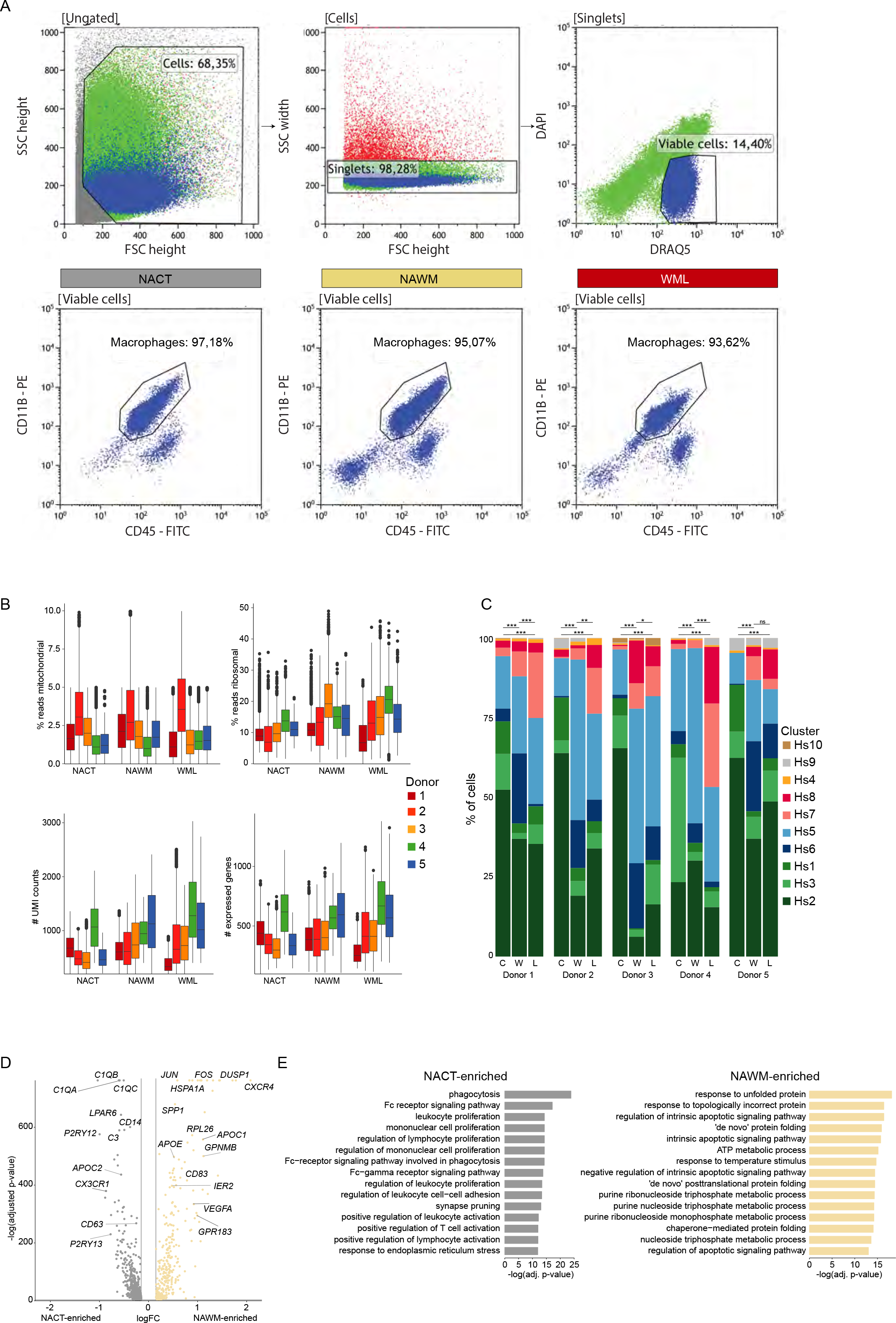
Supplementary data related to figure 3. (**A**) Representative plots depicting FACS strategy to obtain live CD45^pos^CD11B^pos^ cells. (**B**) Boxplots per sample depicting several indicated quality control parameters per cell. (**C**) Stacked bar plots depicting cluster distribution per sample with statistical analysis (Chi-squared test). C = NACT; W = NAWM; L = WML. (**D**) Volcano plot depicting significantly differentially expressed genes between NAWM and NACT samples (paired design). (**E**) Bar plots depicting top 15 gene ontology terms associated with DEGs between NACT and NAWM cells. *: p < 0.05; **: p < 0.01; ***: p <0.001. Abbreviations: NACT = normal appearing cortical tissue; NAWM = normal appearing white matter; WML = white matter lesion.

**Supplementary Figure S3.**
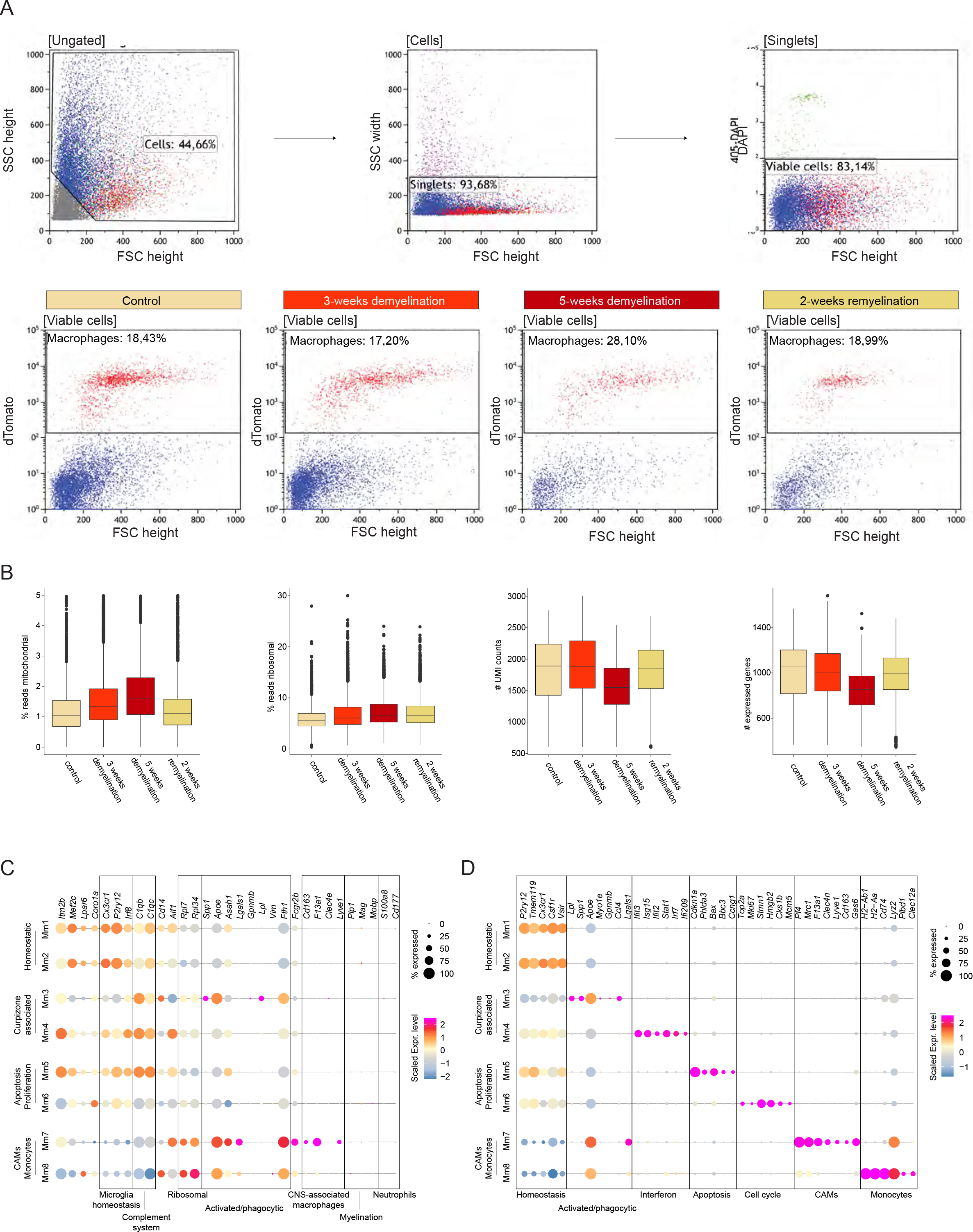
Supplementary data related to figure 4. (**A**) Representative plots depicting FACS strategy to obtain live Cx3cr1^pos^ cells. (**B**) Boxplots per sample depicting several QC indicated stats per cell. (**C**) Dot plot depicting expression of mouse homologs of the genes depicted in figure 3F. (**D**) Dot plot depicting expression of marker genes of each mouse scRNAseq cluster. Size of the symbols depicts the fraction of cells expression the gene, color scale depicts average expression level. Abbreviations: CAM = CNS-associated macrophages.

**Supplementary Figure S4.**
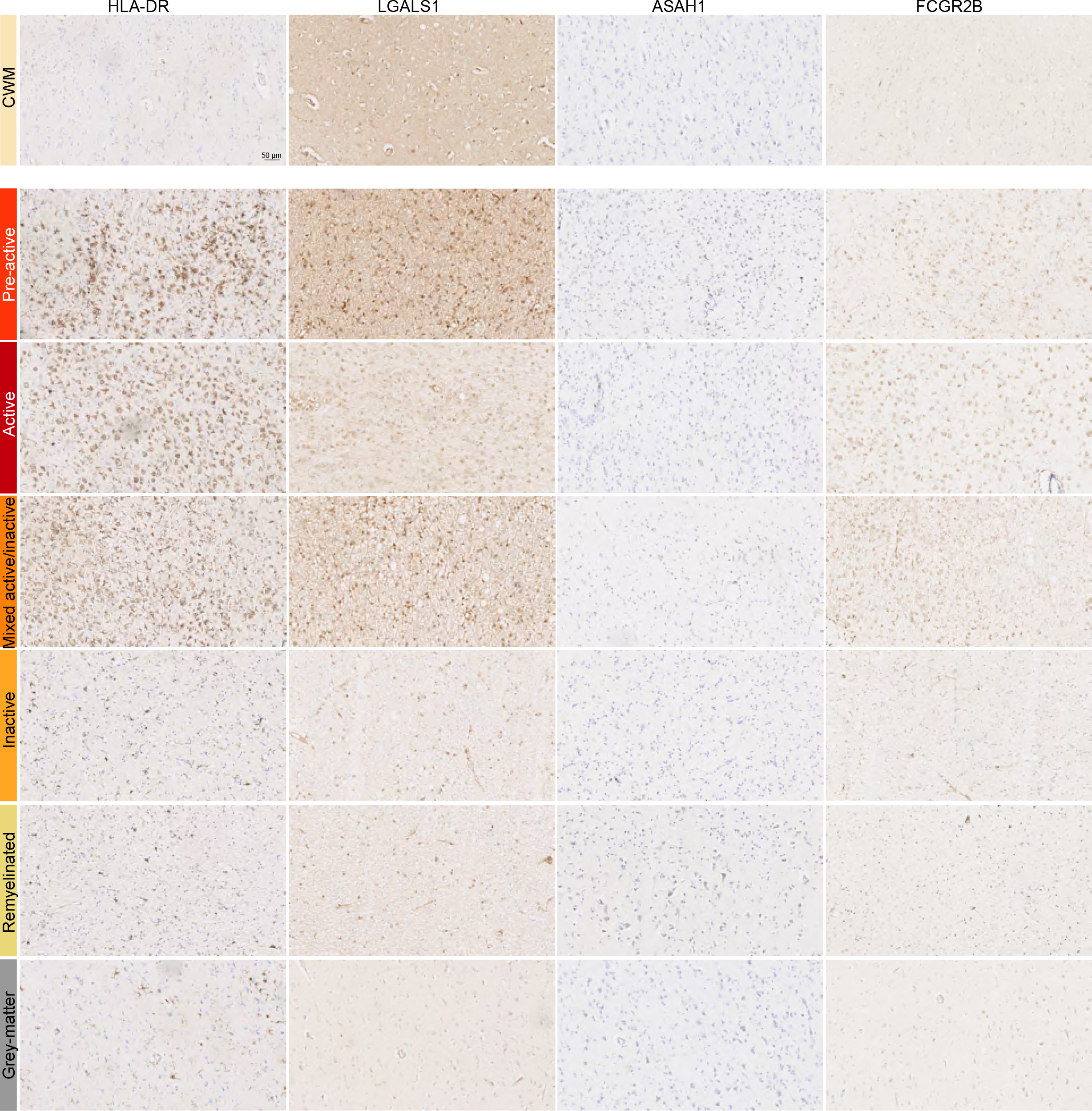
Supplementary data related to figure 5. Representative images of immunohistochemistry for HLA-DR, LGALS1, ASAH1 and FCGR2B in CTR donors and 6 lesion types in MS donors. Abbreviations: CWM = control white matter (CTR donor).

